# Sampling native-like structures of RNA-protein complexes through Rosetta folding and docking

**DOI:** 10.1101/339374

**Authors:** Kalli Kappel, Rhiju Das

## Abstract

RNA-protein complexes underlie numerous cellular processes including translation, splicing, and posttranscriptional regulation of gene expression. The structures of these complexes are crucial to their functions but often elude high-resolution structure determination. Computational methods are needed that can integrate low-resolution data for RNA-protein complexes while modeling *de novo* the large conformational changes of RNA components upon complex formation. To address this challenge, we describe a Rosetta method called *RNP-denovo* to simultaneously fold and dock RNA to a protein surface. On a benchmark set of structurally diverse RNA-protein complexes that are not solvable with prior strategies, this fold-and-dock method consistently sampled native-like structures with better than nucleotide resolution. We revisited three past blind modeling challenges in which previous methods gave poor results: human telomerase, an RNA methyltransferase with a ribosomal RNA domain, and the spliceosome. When coupled with the same sparse FRET, cross-linking, and functional data used in previous work, *RNP-denovo* gave models with significantly improved accuracy. These results open a route to computationally modeling global folds of RNA-protein complexes from low-resolution data.

## Introduction

RNA-protein interactions underlie many critical cellular processes from translation, splicing, and telomere extension to regulation of mRNA stability, alternative splicing, and subcellular localization (Gerstberger et al., 2014; Mitchell and Parker, 2014). Many of these roles require the formation of intricate three-dimensional structures, but the structural heterogeneity and transient nature of many RNP states render them invisible to all but the lowest resolution methods, such as FRET, crosslinking, and mutagenesis tests. For such states, computational techniques will be needed for “hybrid” structure determination integrating low-resolution data into structural models (Schlundt et al., 2017; Ward et al., 2013). Such strategies have proved useful for RNA and proteins separately (Miao et al., 2017; Moult et al., 2017; Ward et al., 2013; Weinreb et al., 2016), but they are not yet in widespread use for RNA-protein complexes because the necessary computational structure prediction methods have not yet been developed.

The majority of existing structure prediction tools for RNA-protein complexes focus on rigid-body docking of RNA and protein partners (Tuszynska et al., 2014). These methods have achieved impressive success when predicting structures of complexes from the corresponding bound RNA and protein structures (Huang and Zou, 2014; Huang et al., 2013; Li et al., 2012; Setny and Zacharias, 2011). However, they typically perform poorly in the more realistic case of starting from the unbound RNA and protein structures (Lensink and Wodak, 2010). This limitation is largely due to the flexibility and conformational variability of RNA; protein-bound RNA structures often differ considerably from the corresponding unbound RNA structures (Rau et al., 2012).

To help address this challenge, a fragment-based method for predicting structures of single-stranded RNA bound to proteins was recently developed (Chauvot de Beauchene et al., 2016; Chauvot de Beauchene, 2016). Starting from a protein structure and the positions of a few anchor RNA nucleotides, this method was able to predict structures of RNA recognition motif (RRM) and Puf proteins bound to single-stranded RNA with high accuracy. However, this method neglects intramolecular RNA interactions and assumes that every RNA nucleotide interacts with the protein, making it applicable to only a small subset of RNA-protein complexes. Currently, there are no computational tools that can model arbitrary protein-bound RNA structures *de novo*, though in principle this could be accomplished by combining RNA structure prediction (“folding”) and RNA-protein docking methods sequentially in a “prefold-then-dock” strategy.

Here, we present tests of this prefold-then-dock strategy on a benchmark set of ten diverse RNA-protein complexes and find that it does not lead to accurate models, suggesting that a new modeling approach is needed. We then describe a method *RNP-denovo* to model RNA-protein complexes by simultaneously folding and docking RNA to a protein surface. This fold-and-dock approach is implemented in Rosetta and combines the FARNA method for RNA folding (Cheng et al., 2015) with RNA-protein binding and a statistical RNA-protein score function. For the benchmark set of ten RNA-protein complexes, starting from the unbound protein structure and a few RNA residues fixed relative to the protein (to simulate sparse experimental data), *RNP-denovo* recovered native-like models with an average RMSD over the best models of 4.3 Å, which is comparable to what has previously been achieved for low-resolution protein and RNA structure prediction on similar size problems (Das and Baker, 2007; Simons et al., 1997). Additional tests demonstrate the importance of including both RNA-protein and intramolecular RNA interactions when modeling protein-bound RNA structures. Finally, we apply our *RNP-denovo* fold-and-dock method to previous structure modeling challenges based on limited experimental data for the human telomerase core RNP, an RNA methyltransferase, and the human spliceosomal C complex active site. We find improvements over previous blind models in all cases, and achieve the correct global folds in the telomerase and RNA methyltransferase cases. These results demonstrate the applicability of this method to real modeling challenges and highlight areas for future improvement. Accurate fully *de novo* prediction of protein-bound RNA structures is not yet feasible, but we expect the method described here to be immediately useful for modeling arbitrary RNA-protein complexes in cases where sparse experimental data are available.

## Results

### Testing a prefold-then-dock approach

To evaluate different protocols for modeling RNA-protein complexes, we built models for a set of ten RNA-protein complexes for which crystal structures are available. The benchmark set contains relatively small RNA-protein complexes with between 94 and 417 protein residues and between 7 and 45 RNA nucleotides per system. This size range is typical for initial structure prediction tests for RNA alone, proteins alone, and protein-protein docking (Das and Baker, 2007; Gray et al., 2003; Simons et al., 1997); tests on larger RNA-protein complexes are described below. Given previous results for low-resolution modeling of RNA and proteins separately, the target modeling accuracy for systems in this size range is around 2-7 Å RMSD (Das and Baker, 2007; Simons et al., 1997). Because the methods considered here do not include final full-atom refinement, we focused on evaluating whether native-like conformations are sampled. This is important because current high-resolution refinement methods typically do not modify structures dramatically. We therefore report the best RMSD accuracy of the top 100 scoring models (out of thousands) in all cases. This procedure is also consistent with typical evaluation criteria in structure prediction challenges, where multiple models are often considered and the number of models is increased to assess progress on more difficult problems with large search spaces (Lensink et al., 2017; Miao et al., 2017; Miao and Westhof, 2017; Moult et al., 2018).

To first address whether a combination of existing tools might be sufficient to predict structures of protein-bound RNA, we tested a prefold-then-dock strategy on a set of ten RNA-protein complexes using the FARFAR method (Cheng et al., 2015) to fold the RNA and then RPDock to dock the resulting RNA structures (Huang et al., 2013). We first tested whether RPDock could accurately predict structures of RNA-protein complexes if starting from the bound protein structure and bound RNA structure, i.e., if the RNA structure could somehow be predicted perfectly. Indeed, RPDock recovered near-native models with RMSDs ≤ 1.5 Å for the best of the top 100 scoring models in nine out of ten cases, with mean RMSD of 1.8 Å over all ten cases. However, if we use the unbound protein structures, as is more realistic for macromolecule docking, the mean RMSD overall increases to 7.1 Å. Furthermore, the results became significantly worse if we assume that the bound RNA structures are unknown, as is typically the case in realistic modeling scenarios. For these tests, we folded the RNA with FARFAR and clustered the resulting RNA structures, retaining the centers of the ten most populated clusters for subsequent docking in the hopes of capturing some of the conformational heterogeneity of the unbound RNA and including conformations similar to the bound structures. After docking these structures with RPDock, the mean RMSD increased to 13.8 Å, with RMSDs worse than 11 Å in nine out of ten cases (Figure 1A-K, Figure S1A, see Methods for complete details). Indeed even assuming the best possible docking by aligning predicted RNA structures to the native RNA coordinates, the mean RMSD remained at 6.5 Å, with RMSDs > 7 Å in five out of ten cases (Figure S1A). We emphasize that the RMSD values here and below are computed for the best of 100 models and therefore represent ‘best-case’ assessments. The poor results suggest that sequentially folding and docking RNA structures does not generally lead to accurate models of RNA-protein complexes.

**Figure 1.**
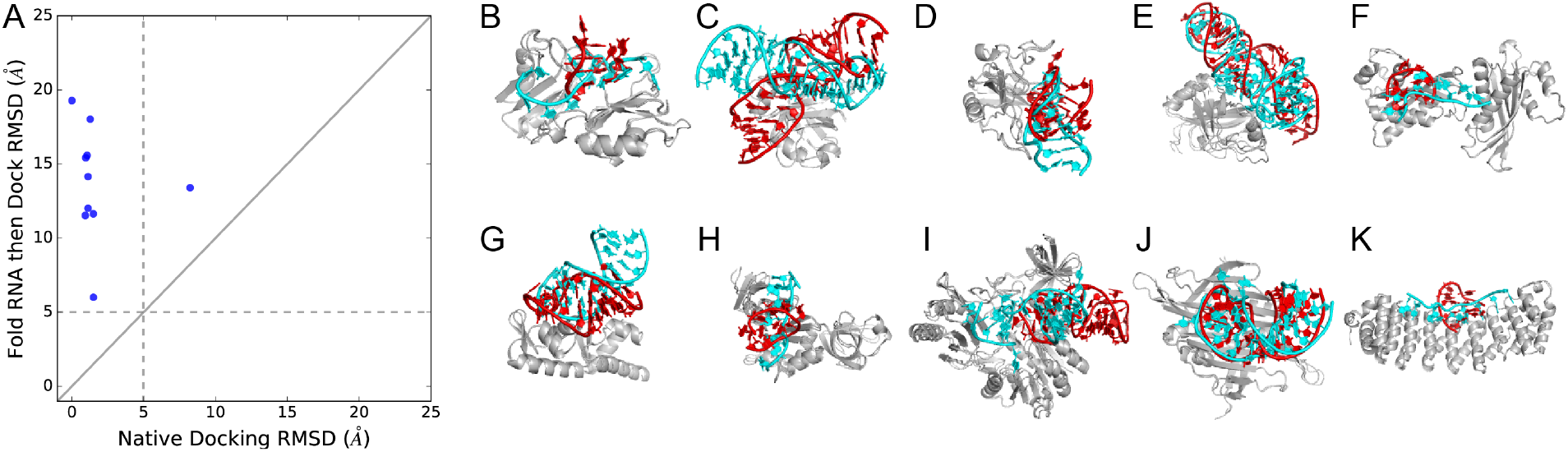
Tests of the best accuracies achievable by a prefold-then-dock strategy. A) The best RMSDs achieved for ten systems for the prefold-then-dock strategy versus the (unrealistic) best case of rigid-body docking of the native RNA to the native protein structures. (B-K) Native structures (RNA in cyan) overlaid with the best models from the prefold-then-dock method (RNA in red, protein in gray) for (B) 1B7F (C) 1DFU, (D) 1JBS, (E) 1P6V, (F) 1WPU, (G) 1WSU, (H) 2ASB, (I) 2BH2, (J) 2QUX, and (K) 3BX2. See also Figure S1.

However, in many realistic cases, the structure prediction problem may be simplified by the availability of limited experimental data. In favorable cases, these data can elucidate a few specific RNA protein contacts and/or relative orientations of the RNA and protein partners. To test the prefold-then-dock strategy in such scenarios, we simulated the availability of limited experimental data by assuming the bound conformations of the 3’ and 5’ nucleotides for the single-stranded RNA-protein complexes (analogous to previous work (Chauvot de Beauchene, 2016)), or the positions of RNA helices relative to the protein for the remaining complexes (see Methods for complete details). When the RNA was folded with these constraints, the mean RMSD was 5.4 Å over the ten RNA-protein complexes, representing an improvement over the naïve prefold-then-dock tests, but still far from the 1.8 Å mean RMSD achieved when docking the bound RNA conformations (Table 1, Figure S1B). Together these results suggest that a prefold-then-dock strategy alone is insufficient to recover near-native conformations of RNA-protein complexes, and that a strategy that allows simultaneous optimization of the RNA fold and the docked conformation may improve predictions.

**Table 1.** Accuracy of prefold-then-dock versus Rosetta *RNP-denovo* fold-and-dock.

| System | Best RMSD of top 100 scoring models (Å) |  |
| --- | --- | --- |
|  | Prefold-then-dock<sup>1</sup> | Rosetta *RNP-denovo* fold-and-dock |
| Sex-lethal RRM (1b7f) | 6.7 | 4.2 |
| Ribotoxin restrictocin – SRL analog (1jbs) | 3.1 | 3.0 |
| HutP antitermination complex (1wpu) | 7.3 | 3.9 |
| mRNA binding domain of SelB elongation factor (1wsu) | 2.4 | 2.4 |
| NusA transcriptional regulator (2asb) | 5.6 | 3.1 |
| Methyltransferase RumA in complex with rRNA (2bh2) | 6.5 | 5.8 |
| PP7 coat protein and viral RNA (2qux) | 5.2 | 4.7 |
| Puf4 bound to 3' UTR of target transcript (3bx2) | 6.3 | 3.8 |
| SmpB-tmRNA complex (1p6v) | 5.2 | 6.3 |
| *E. coli* L25-5S rRNA (1dfu) | 5.2 | 5.5 |
| Human telomerase<sup>2</sup> | 51.4 (25.5)<sup>3</sup> | 15.0 |
| RNA methyltransferase (CAPRI T33) | 18.2 (31.0)<sup>4</sup> | 13.6 |
| Human spliceosome | 13.8 (34.5)<sup>5</sup> | 8.0 |
| **Average** | **10.5** | **6.1** |

<sup>1</sup>For the ten small RNA-protein systems, the RNA was folded keeping the same residues fixed relative to the protein as in the *RNP-denovo* fold-and-dock runs (therefore, docking was not performed; see Figure 1 and Figure S1 for full prefold-then-dock results).
<sup>2</sup>Results for models built without FRET data. RMSDs calculated over pseudoknot ends (see Methods). Accuracy of best of top 100 previous blind models shown in parentheses.
<sup>3</sup>Because the position of the template RNA relative to the protein is known, prefold-then-dock was not attempted. Instead, the results here are for the *RNP-denovo* fold-and-dock runs with just RNA-only score terms and RNA-protein sterics. Accuracy of best of top 100 previous blind models shown in parentheses.
<sup>4</sup>Accuracy of the best of the previously submitted blind models shown in parentheses.
<sup>5</sup>Accuracy of the previously published blind model shown in parentheses. RMSDs were calculated over active site residues (see Methods).

### Developing a fold-and-dock method

Motivated by these results, we hypothesize that folding the RNA in the context of the protein rather than pre-folding and docking would further improve the accuracy of computational models. We developed a fold-and-dock algorithm *RNP-denovo* in Rosetta by modifying the FARNA algorithm for RNA folding to include RNA-protein docking moves and to take into account RNA-protein interactions. Currently, *RNP-denovo* does not include full-atom refinement; as decribed above, we wish to cleanly test here whether native-like conformations are sampled with a low-resolution protocol. This sampling is important because current high-resolution refinement methods typically do not modify structures dramatically. We also developed a new RNA-protein statistical potential to score conformations of RNA as it makes contact with a protein surface. While prior studies have developed statistical potentials for RNA-protein docking, we sought a scoring function that could be rapidly computed at the same time as Rosetta RNA score terms and have a similar level of coarse graining. Overall, the new scoring function includes all previously published score terms describing RNA structure (Das and Baker, 2007), as well as new terms describing interactions between RNA and proteins. As with the Rosetta RNA statistical potential, we took a coarse-grained knowledge-based approach. Score terms were based on the frequencies of interactions observed in a non-redundant set of 154 crystal structures of RNA-protein complexes with resolution better than 2.5 Å, curated from the Protein Data Bank (PDB).

The score terms describe three major features of RNA-protein interactions. First, pseudo pairs between nucleotides and protein sidechains have been observed and are thought to contribute to both the specificity and affinity of RNA-protein interactions (Kondo and Westhof, 2011). To capture these effects, we developed a potential based on the distributions of protein sidechain centroids in the plane of each of the four RNA bases. As described previously for the RNA score terms (Das and Baker, 2007), a coordinate system was set up on each base with the origin at the centroid of the base heavy atoms, the x-axis going through the N1 atom for purines or the N3 atom for pyrimidines, and the z-axis perpendicular to the plane of the base. Analysis of the protein sidechain distributions indeed revealed positional preferences within the plane of the base (Figure 2A, Figure S2A), though the interactions were not as highly stereotyped as for example, RNA base pairs (Figure 2B, C). Statistical potentials were derived by taking the negative logarithm of these frequencies (see Methods). These terms include RNA-protein pseudo pairs previously identified by expert inspection (Kondo and Westhof, 2011) as well as less stereotyped RNA-protein contacts.

**Figure 2.**
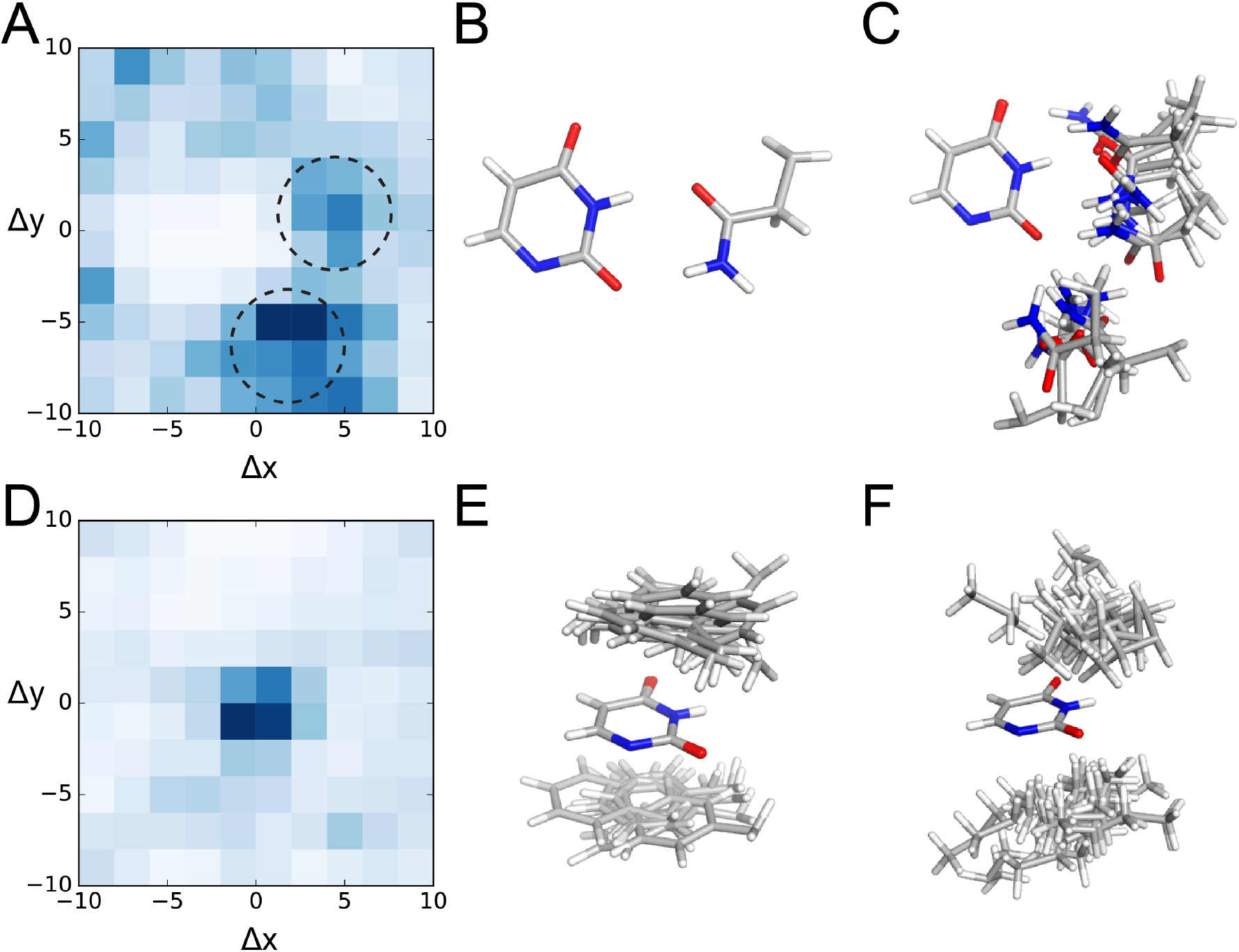
Statistical RNA-protein potential in Rosetta. A) The distribution of glutamine sidechain centroids around uracil in the plane of the base (0 < |z| < 3 Å), from the non-redundant set of RNA-protein crystal structures from the PDB (darker blue represents higher frequency). Distributions of all protein sidechains around all four RNA bases are shown in Figure S2A. B) A pseudo-pair between glutamine and uracil. C) Conformations from the two major hotspots circled in (A) show that the interactions between glutamine and uracil are not highly stereotyped. D) The distribution of phenylalanine sidechain centroids around uracil above and below the plane of the base (3 < |z| < 6.5 Å; darker blue represents higher frequency). Distributions of all protein sidechains around all four RNA bases are shown in Figure S2B. E) Representative conformations from the hotspot in (D) show stereotyped stacking interactions. F) Conformations of valine around uracil also reveal frequent stacking interactions. See also Figures S2, S3, S4.

Second, the potential captures the effect of stacking interactions frequently found at RNA-protein interfaces (Rahman et al., 2015). Analysis of the distributions of protein sidechains above and below the plane of the base revealed that the aromatic amino acids tryptophan, tyrosine, and phenylalanine and two additional hydrophobic amino acids, valine and leucine, frequently stack on RNA bases (Figure 2D-F, Figure S2B). A stacking bonus is encoded in the potential for any of these five sidechains with 3.0 Å < |z| < 6.5 Å and 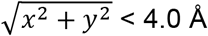, again analogous to the bonus for RNA-RNA stacking.

Third, the potential includes the effect of interactions with the RNA backbone, which often confer both affinity and structural specificity to RNA-protein interactions (Iwakiri et al., 2012). Scores were inferred by taking the negative logarithm of the frequencies of distances between RNA phosphate atoms and protein sidechain centroids (see Methods). The resulting statistical potentials exhibit several expected features (Figure S3A), most notably that interactions with positively charged amino acids (arginine and lysine) are among those that score most favorably and interactions with polar amino acids are generally preferred over interactions with nonpolar amino acids.

To account for additional interactions that may be missed by the three score terms described above, we also included a general distance dependent potential based on the observed distributions of distances between representative RNA and protein atoms (Figure S3B), analogous to several prior efforts (Guilhot-Gaudeffroy et al., 2014; Huang and Zou, 2014; Li et al., 2012; Perez-Cano et al., 2010; Setny and Zacharias, 2011; Simons et al., 1997; Tuszynska and Bujnicki, 2011; Zheng et al., 2007). Finally, steric clashes between the RNA and protein are penalized in a manner similar to clashes within the RNA (Das and Baker, 2007). A penalty is applied when representative RNA and protein atoms come within a distance smaller than the minimum distance observed in the set of crystal structures from the PDB (Figure S4).

### Benchmarking RNP-denovo on ten RNA-protein complexes

We benchmarked the performance of the *RNP-denovo* fold-and-dock method on the same set of ten RNA-protein complexes that was used for the prefold-then-dock tests described above. Again, to simplify the problem and simulate the availability of limited experimental data, we assumed the positions of the 5’ and 3’ nucleotides relative to the protein for the singlestranded RNA binding proteins, and the relative positions of the RNA helices for the remaining systems (see Methods). We then built models of the remaining RNA residues in the presence of the protein with *RNP-denovo*. To assess the effect of the RNA-protein score terms, we first built a set of models with a score function that included only the RNA-specific score terms and the RNA-protein steric penalty. The best of the top 100 scoring models (top 1.4%) achieved an average RMSD of 6.0 Å, with RMSDs better than 5 Å in three out of ten cases (Table S1). For models built with all of the RNA-protein score terms included, the average RMSD over the best of the top 100 scoring models improved to 4.3 Å, with RMSDs better than 5 Å in seven out of ten cases (Table 1, Table S1), recovering near-native RNA folds for all systems (Figure 3A-J). In all cases, the inclusion of the RNA-protein score terms resulted in a shift in the distribution of *RNP-denovo* models towards lower RMSDs (Figure 3A-J).

**Figure 3.**
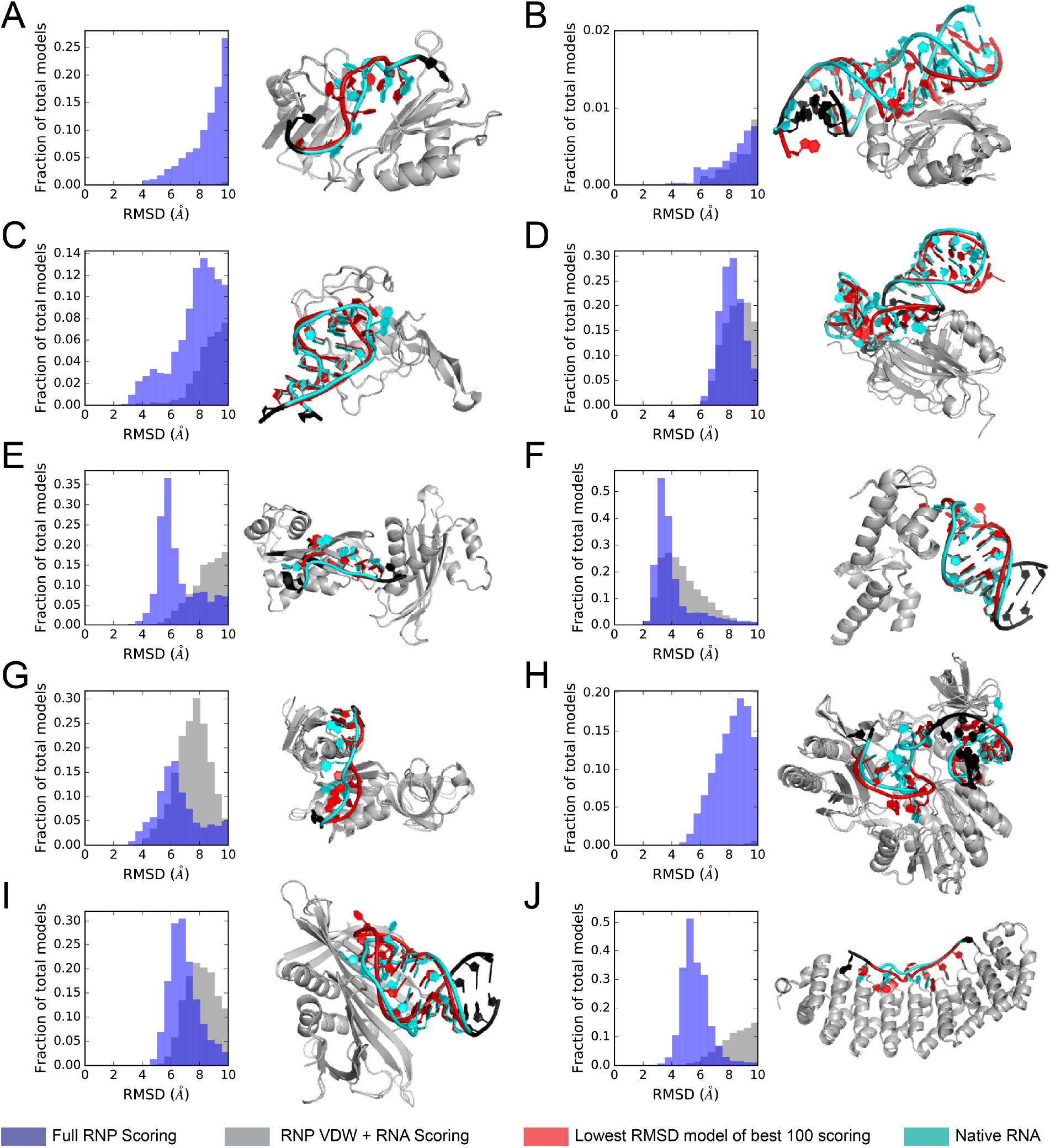
Results of Rosetta *RNP-denovo* folding. (Left) Histograms of *RNP-denovo* RMSDs relative to native for Rosetta models built with the full RNP score function (blue) and the RNP van der Waals term plus RNA-only score terms (gray), and (Right) the best models (by RMSD) out of the top 100 scoring (RNA in red) overlaid with the native RNA-protein complexes (RNA in cyan) for (A) 1B7F, (B) 1DFU, (C) 1JBS, (D) 1P6V, (E) 1WPU, (F) 1WSU, (G) 2ASB, (H) 2BH2, (I) 2QUX, and (J) 3BX2. RNA residues that were kept fixed relative to the protein are colored black. Unbound protein structures (gray) were used for modeling to simulate a realistic prediction scenario. See also Table S1.

### Testing alternative score functions

The major difference between the Rosetta RNA-protein potential described here and previously developed RNA-protein docking potentials is the inclusion of terms describing RNA-RNA interactions, which can safely be neglected for rigid-body docking problems. To test whether existing docking potentials might produce similar results if integrated into an algorithm for structure prediction of RNA at protein interfaces, such as the *RNP-denovo* fold-and-dock method described here, we rescored our *RNP-denovo* models with the 3dRPC (Huang et al., 2016) and DARS-RNP docking potentials (Tuszynska and Bujnicki, 2011). For the singlestranded RNA binding proteins, the docking potentials picked out models with accuracy similar to the full Rosetta RNP potential (Table S2, Figure 4A). However, for three of the RNA-protein systems containing structured RNA, the docking potentials picked out models that deviated significantly from the native conformations with RMSDs ≥ 14.7 Å. Over all ten systems, the average RMSD of the best scoring models was 11.6 Å for 3dRPC and 10.2 Å for DARS-RNP compared to 6.4 Å for the Rosetta RNA-protein score function (Table S2, Figure 4A). The docking potentials performed worst for 1DFU, with the distribution of model accuracy over the top scoring models shifting dramatically towards poorer RMSDs (Figure 4B). The best scoring *RNP-denovo* model picked out by both DARS-RNP and 3dRPC adopts a conformation in which the two RNA strands, which interact in the native complex, wrap around opposite sides of the protein to maximize the number of RNA-protein contacts (Figure 4C). As an additional comparison, we rescored the *RNP-denovo* models with just the five Rosetta RNA-protein (intermolecular) score terms. The results were similar to the poor results with prior RNA-protein docking potentials, with an average RMSD over the best scoring models for each of the ten systems of 10.5 Å (Table S2, Figure S5). These results highlight the importance of RNA-RNA interactions in RNA-protein complexes. All “decoy” models are being made available at https://purl.stanford.edu/gt072md8147 to allow testing of new RNP scoring functions.

**Figure 4.**
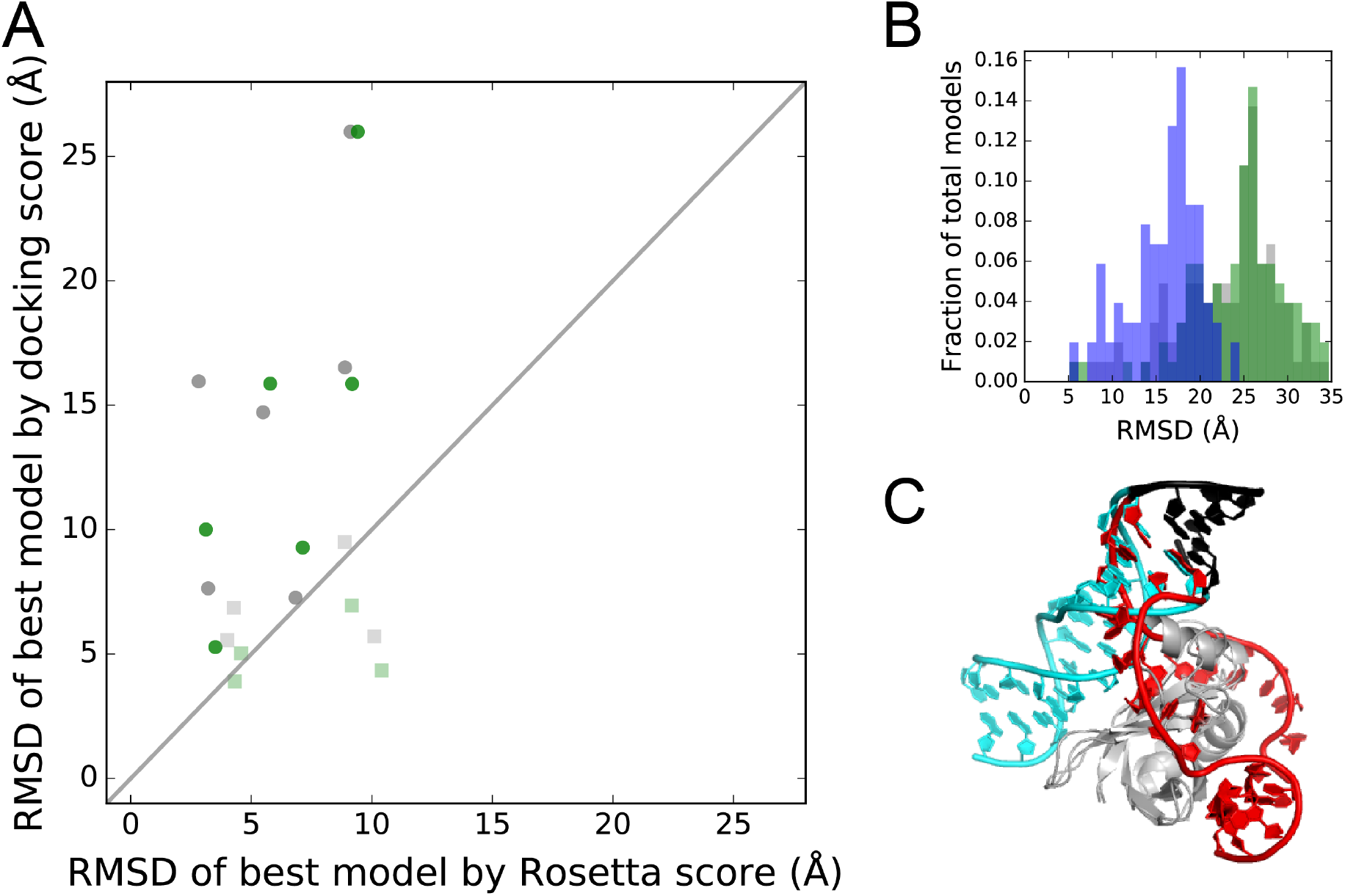
Rescoring *RNP-denovo* Rosetta models with docking potentials. A) The RMSD of the best scoring Rosetta models assessed by the DARS-RNP (green) and 3dRPC (gray) docking potentials versus by the Rosetta RNP score function for systems shown in Figure 3. Values for single-stranded RNA binding proteins are shown as squares. B) Distributions of RMSDs for the top 100 scoring models assessed by the DARS-RNP (green), 3dRPC (gray), and Rosetta RNP potentials (blue). C) The best scoring model assessed by 3dRPC for 1dfu (RNA colored red) overlaid with the native structure (RNA colored cyan). Residues that were kept fixed during modeling are colored black. See also Figure S5 and Table S2.

### Applying RNP-denovo to three past modeling challenges of large RNPs

To test whether *RNP-denovo* would be useful for real modeling problems, we revisited three past modeling challenges for large RNA-protein complexes in which only sparse experimental data were available. These three systems are substantially larger than the complexes in our initial benchmark set; the average number of protein and RNA residues per system is 702 and 117 residues, respectively, compared to 215 and 21 residues, respectively, for the initial benchmark set. Due to the increased size of these problems, we expected the RMSD values to be higher than for the initial benchmark set. Using a previously determined relationship between number of residues and RMSD (Carugo and Pongor, 2001), we calculated that the structural similarity specified by an RMSD of 4.3 Å (average for the initial benchmark set) for complexes with 21 RNA residues on average would correspond approximately to an RMSD of 21 Å for the larger complexes (see Methods). We therefore targeted RMSD accuracies of better than 21 Å as representative of correct global fold recovery for these three larger systems. First, we built models of the human telomerase core RNP based on FRET measurements, which provided ten distance restraints between specific pairs of RNA residues (Parks et al., 2017). Models of this system were previously built in 2015 using FARNA with these FRET constraints and an additional score term that penalized RNA-protein steric clashes (Parks et al., 2017). Here, we followed the previous modeling procedure, but used the *RNP-denovo* fold-and-dock method and the more advanced Rosetta RNP statistical potential described here (see Methods). As before, the positions of the template hybrid and CR4/5 RNA were kept fixed relative to the homology model of the telomerase reverse transcriptase protein (TERT). Although a high-resolution structure of the human telomerase core RNP has still not been determined, the accuracy of the newly and previously built models could be assessed by comparing to the recently solved 7.7 Å resolution cryoEM structure of the human telomerase RNP (Nguyen et al., 2018) (which was determined after the original and *RNP-denovo* models were built). Specifically, we considered the positioning of the highly conserved RNA pseudoknot motif relative to the TERT protein. Qualitatively, both the previously built and the new *RNP-denovo* fold-and-dock models agree well with the cryoEM structure, with the pseudoknot positioned on the correct face of TERT. The best *RNP-denovo* model (out of the top 100 scoring models) has an improved RMSD accuracy of 13.2 Å over the ends of the pseudoknot motif (see Methods), compared to 17.1 Å for the best of the top 100 previously built models (Figure S6A, S8B, S8C). Although the accuracy here is worse than for the systems in the initial benchmark set, it is still reasonable given the increased size of this problem. Notably, at 13.2 Å RMSD, the global fold is still correctly recovered (Figure 5A, middle and right panels).

**Figure 5.**
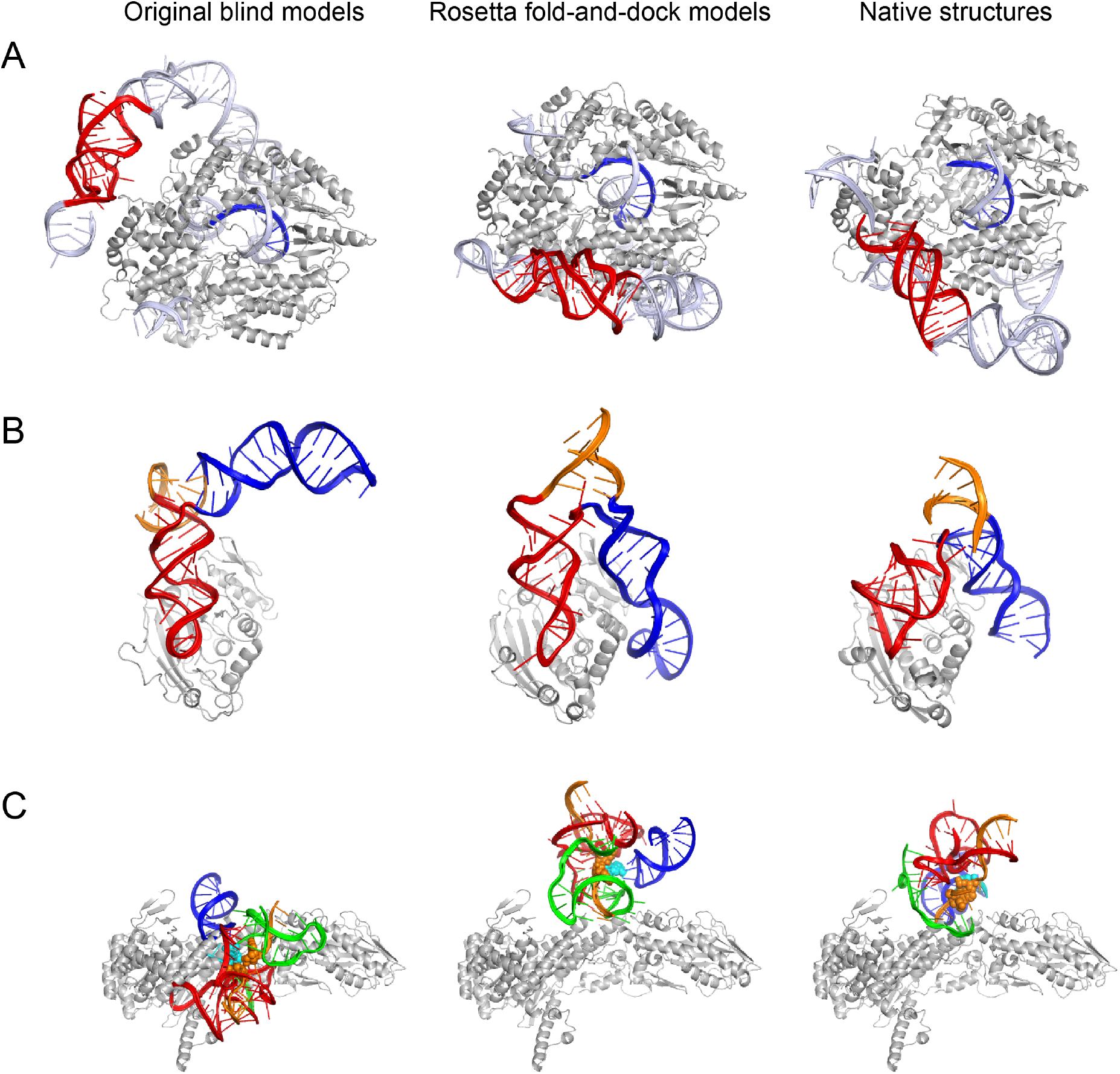
Revisiting three past modeling challenges with the Rosetta *RNP-denovo* fold-and-dock method. (A) The best of the previously selected blind human telomerase core RNP models built without FRET data (Parks et al., 2017) (left; RMSD over select pseudoknot residues = 78.8 Å; pseudoknot RNA motif colored red, template RNA colored blue, other modeled RNA colored light blue, protein colored gray), the best *RNP-denovo* fold-and-dock model by RMSD of the top 100 scoring models built without FRET (middle; RMSD over select pseudoknot residues = 15.0 Å), and the cryoEM structure of human telomerase (Nguyen et al., 2018) (right). (B) The best of the previously submitted RNA methyltransferase CAPRI T33 models (left; RMSD = 31.0 Å; RNA colored blue, red, and orange; protein colored gray), the best of the top 100 scoring *RNP-denovo* fold-and-dock models (middle; RMSD = 13.6 Å), and the best T34 model, which closely resembles the crystal structure (right; interface RMSD to crystal structure = 1.5 Å). (C) The previously published human spliceosomal C complex model (Anokhina et al., 2013) (left; RMSD over key active site residues (shown as spheres) = 34.5 Å, U2 RNA colored green, U5 RNA colored blue, U6 RNA colored red, intron colored orange, 5’ exon colored cyan, protein colored gray), the best RMSD model of the top 100 scoring *RNP-denovo* fold-and-dock models of the human C complex (middle; RMSD over key active site residues = 8.0 Å), and the cryoEM structure of the yeast C complex (right). See also Figure S6.

We additionally tested whether the inclusion of the FRET data was necessary to build accurate models of the telomerase core RNP, as was noted for the previously built models (Parks et al., 2017). The top scoring models built without FRET data using *RNP-denovo* were on average more accurate than top scoring models built with the previous method without FRET data (Figure S6D, S8E). The most accurate *RNP-denovo* model of the top 100 scoring had an RMSD of 15.0 Å, compared to 51.4 Å for models built with the previous method (scoring with RNA score terms and RNA-protein sterics only) and 25.5 Å for the top 100 scoring previously built blind models (Figure 5A). Additionally, predicted FRET values calculated from models built using the previous method correlate poorly with the experimental FRET values, with a maximum correlation of 0.37 (Figure S6F). This correlation is improved for many of the models built with the new *RNP-denovo* method, with the best RMSD model achieving the maximum correlation of 0.60 (Figure S6F). These results suggest that in contrast to the previous approach, *RNP-denovo* does not require FRET data to sample accurate models, though inclusion of the data still enriches for and helps select accurate models.

We then revisited Target 33/34, an RNA methyltransferase, from the 2008 CAPRI blind modeling challenge (Janin, 2010). Target 33 challenged modelers to build a structure of the full RNA-protein complex, starting from sequence only. Subsequently, for Target 34 modelers were provided with the crystal structure of the bound RNA and asked to predict the structure of the RNA-protein complex. Originally, models for Target 33 were built with Rosetta using an *ad hoc* prefold-then-dock approach, with restraints based on RNA chemical structure probing data, highly conserved protein residues likely to interact with the RNA, and a homology model of the methyltransferase ((Fleishman et al., 2010) and Methods). Although the crystal structure of the complex has not yet been released, the accuracy of the models can be assessed by comparing to the best Target 34 model of the bound RNA crystal structure docked to the protein, which CAPRI evaluators confirmed to be close to the crystal structure of the complex (interface RMSD (I_rmsdBB) of 1.5 Å; http://www.ebi.ac.uk/msd-srv/capri/round15/round15.html). Using this structure for comparison, the previously submitted Target 33 model achieved 31.0 Å RMSD over the RNA after aligning over the protein. To determine whether the *RNP-denovo* fold-and-dock method could build more accurate models, we applied it to this problem and additionally repeated the prefold-then-dock modeling to control for possible changes to the RNA modeling procedure (see Methods). When comparing the accuracy over the full complex (RMSD of the RNA after aligning to the protein), the best of the top 100 scoring *RNP-denovo* models was more accurate than the best of the top 100 scoring prefold-then-dock models, with RMSDs of 13.6 Å and 18.2 Å, respectively (Figure S6G, Figure 5B).

Finally, we applied the *RNP-denovo* fold-and-dock method to build a model of the human spliceosomal C complex active site. Prior to the determination of high-resolution cryoEM structures, Anokhina et al. built a model of the core RNA elements bound to the Prp8 protein based on RNA chemical structure probing data, crosslinking data, homology to the group II intron, and a crystal structure of the Prp8 protein alone (Anokhina et al., 2013). While many features of this model, particularly the relative arrangement of the intron and U2 and U6 RNA, agree well with the later solved cryoEM structures, the positioning of these RNA elements relative to the Prp8 protein was not highly accurate with 34.5 Å RMSD over key active site RNA residues (the 3’ residue of the 5’ exon, the 5’ residue of the intron, and the residue immediately 3’ of the branchpoint A) after alignment over Prp8 (Figure 5C). This inaccuracy can potentially be explained by the fact that the RNA model was docked rigidly to Prp8 despite explicitly noted uncertainty in the U5 RNA positioning (Figure S6H, I). Applying the Rosetta prefold-then-dock approach to this problem gave improved RMSD accuracy over active site residues of 13.8 Å for the best of the top 100 scoring models for which the crosslinking constraints were satisfied, although the global fold over the entire RNA-protein complex was not recovered (Figure S6I). The complete *RNP-denovo* fold-and-dock method has the potential to further improve the accuracy of the model by explicitly accounting for the uncertainty in U5 RNA positioning and allowing it to move relative to the rest of the RNA during docking to Prp8. Starting from the published RNA model and using the same crosslinking constraints, but allowing U5 to move, the *RNP-denovo* Rosetta fold-and-dock method resulted in models of the Prp8-RNA complex that were more accurate than the previously published model and the prefold-then-dock models although the global fold was still not completely accurate (Figure 5C). Over just active site residues, the best of the top 100 scoring models for which the crosslinking constraints were satisfied achieved 8.0 Å RMSD (Figure S6I, Figure 5C orange and cyan space-filled residues).

Overall, for three large RNA-protein systems with sparse experimental restraints, the *RNP-denovo* method resulted in models with improved or similar accuracy compared to previously published models. These tests suggest that this method is useful for real modeling challenges, sampling an ensemble of models with biologically correct global folds or placement of functional residues.

## Discussion

Structure prediction of RNA-protein complexes has remained a relatively unexplored area of research, with efforts predominantly focused on RNA structure prediction without consideration of RNA-protein binding or separately on RNA-protein rigid-body docking. A critical bottleneck is the computational difficulty of sampling *de novo* the new conformations that RNAs form when interacting with protein surfaces. Tests presented here show that combining existing tools in a prefold-then-dock strategy does not generally lead to accurate models of RNA-protein complexes, and that simultaneous optimization of RNA structure and rigid-body orientation is necessary to more accurately predict the structures of these complexes. Over a benchmark set of ten RNA-protein complexes, with the assumption of a few RNA-protein contacts, a Rosetta *RNP-denovo* fold-and-dock method recovered native-like RNA folds in all cases. The new knowledge-based RNA-protein potential implemented in Rosetta enriched sampling of near-native models. Additionally, models favored by potentials that do not include intramolecular RNA interactions were less accurate compared to those favored by the full Rosetta RNP potential for systems containing structured RNA, suggesting that it is necessary to balance consideration of both RNA-RNA and RNA-protein interactions to accurately fold protein-bound RNA structures. Finally, application of the *RNP-denovo* fold-and-dock method to three past modeling challenges of large RNA-protein systems resulted in improved accuracy compared to previously published models built with other methods and to prefold-then-dock methods, suggesting that the *RNP-denovo* will be useful in real modeling scenarios. Overall, *RNP-denovo* appears to resolve a critical sampling bottleneck for *de novo* prediction of protein-bound RNA structures. We expect the method to be widely useful for structure determination, particularly because Rosetta already allows integration of numerous kinds of limited experimental data.

Although *RNP-denovo* here represents an advance in our ability to predict the structures of RNA-protein complexes, fully *de novo* structure prediction without experimental data remains an unsolved challenge. The benchmark on the ten small RNA-protein systems presented here relied on having limited information about specific RNA-protein contacts; without this information, we do not yet have the tools to accurately predict structures of RNA-protein complexes. Additionally, the tests on three larger RNA-protein systems suggest that *RNP-denovo* will be useful for real modeling challenges, but also highlight that high-accuracy RNA-protein modeling remains an unsolved problem. Our results suggest several possible reasons as to why RNA-protein structure prediction remains difficult. First, RNA-protein interactions, unlike RNA-RNA base pairing, are not highly stereotyped, making the development of a predictive low-resolution potential difficult. Second, the development of a statistical potential is hindered by the relatively small number of RNA-protein structures in the PDB; our non-redundant set of crystal structures contains only 154 systems. Finally, the overall conformation of an RNA-protein complex is determined by a balance of both intermolecular and intramolecular interactions, resulting in a folding landscape with many local energy minima in which just one of these sets of interactions may be optimized at the expense of the other. This scenario was observed in the tests of 1DFU structure prediction presented here: in addition to near-native models, there were many models generated in which the RNA-protein contacts were maximized at the expense of RNA structure. Efficiently sampling these conformations for large systems and in fully *de novo* tests will be a challenge.

The results described here suggest two additional areas for future improvement. First, the success of this method relies on having limited experimental data. Here for the benchmark of small RNA-protein systems, we simulated this situation by assuming specific RNA-protein contacts, and for the three large RNA-protein tests, we used FRET or crosslinking data, or information about highly conserved residues. However this method could be further generalized to include other types of experimental information such as NMR restraints (Zhang et al., 2018), contacts derived from evolutionary couplings (Weinreb et al., 2016), or SAXS data (Schneidman-Duhovny et al., 2012; Schwieters et al., 2018), as we have accomplished recently for cryoEM data, achieving models with near-atomic accuracy in blind challenges (Kappel et al., 2018). Second, improving the accuracy of this method and discriminating the top model (rather than the top 100) will require new high-resolution refinement methods that can be applied to RNA-protein complexes. Currently, conformations are scored exclusively with a low-resolution knowledge-based potential. Refining these low-resolution structures with a full-atom energy function will likely be necessary to improve the overall accuracy of the models and to improve discrimination of near-native versus non-native conformations.

## Acknowledgements

We thank members of the Das lab for helpful discussions and members of the Rosetta community for sharing code. We thank Juli Feigon and her lab for sharing their coordinates for the 9 Å *Tetrahymena* telomerase cryoEM structure, and Zhichao Miao and Eric Westhof for sharing the coordinates for their human spliceosome model. Calculations were performed on the Stanford BioX^3^ cluster, supported by NIH Shared Instrumentation Grant 1S10RR02664701, and the Sherlock cluster. This work was supported by a Gabilan Stanford Graduate Fellowship (K.K.), a NSF GRFP (K.K.), NIGMS MIRA R35 GM122579 (R.D.), and R01 GM121487 (R.D.).

## Author Contributions

K.K. and R.D. designed the computational approach. K.K. implemented the method and performed the tests and analysis. K.K. and R.D. wrote the manuscript.

## Methods

### The benchmark set of ten RNA-protein complexes

Ten systems were chosen from the non-redundant set of RNA-protein complexes with corresponding unbound protein structures available, described in (Perez-Cano and Fernandez-Recio, 2010). The specific systems were selected manually to represent a diversity of types of RNA-protein interactions (unbound protein structures listed in parentheses): 1DFU (1B75) (Lu and Steitz, 2000; Stoldt et al., 1998), 1B7F (3SXL) (Crowder et al., 1999; Handa et al., 1999), 1JBS (1AQZ) (Yang et al., 2001; Yang and Moffat, 1996), 1P6V (1K8H) (Dong et al., 2002; Gutmann et al., 2003), 1WPU (1WPV) (Kumarevel et al., 2005), 1WSU (1LVA) (Selmer and Su, 2002; Yoshizawa et al., 2005), 2ASB (1K0R) (Beuth et al., 2005; Gopal et al., 2001), 2BH2 (1UWV) (Lee et al., 2004, 2005), 2QUX (2QUD) (Chao et al., 2008), and 3BX2 (3BWT) (Miller et al., 2008). Fixed residues for tests in which some RNA residues were held fixed relative to the protein were selected as follows: for systems containing only single stranded RNA (1B7F, 1WPU, 2ASB, 3BX2), the first and last RNA residues were kept fixed; for 1DFU, 1WSU, and 2QUX, the first three base pairs were held fixed; for 1P6V the first two base pairs of both helices were held fixed (residues 19-20, 37-38, 41-42, 56-57); for 2BH2, the first three base pairs of the RNA helix and the 5’ nucleotide were kept fixed; for 1JBS, the 5’ and 3’ residues were held fixed.

### The prefold-then-dock protocol

For each of the ten structures in the benchmark set, the RNA was folded with the FARFAR method (Cheng et al., 2015). 5000 structures were generated for each system. The resulting structures were clustered in Rosetta, with a clustering radius of 2.0 Å. The centers of the ten most populated clusters were then docked to the unbound protein structures with RPDock (Huang et al., 2013). RMSDs were calculated over RNA heavy atoms after alignment based on the protein coordinates. The best RMSD was selected from the top ten scoring models for each of the ten docked RNA cluster centers (100 structures in total for each system).

To dock the native bound conformations, the RNA and protein chains were extracted from the bound complex, then docked together with RPDOCK. The best RMSD was selected from the top 100 models for each system.

For tests with fixed RNA residues, the RNA residues described above were held fixed during RNA folding in FARFAR. RMSDs were calculated over RNA residues that were not held fixed after alignment to the fixed RNA residues. The best RMSD was selected from the top 100 models for each system (clustering was not performed).

### Assembling a non-redundant set of crystal structures of RNA-protein complexes

All crystal structures containing both RNA and protein chains, with resolution better than 2.5 Å were downloaded from the Protein Data Bank in August 2016. Complexes containing only protein chains of length less than 20 amino acids or RNA chains of length less than four nucleotides were discarded. All protein chains were clustered using blastclust (Altschul et al., 1990) with a 30% sequence identity cutoff using the command “blastclust –i fasta.txt –o blastclust_output.txt –b T –S 30”. Because the protein chains were clustered individually, different protein chains of the same RNA-protein complex could end up in different clusters after this step. This was addressed by merging clusters with members from the same RNA-protein complex. A single representative structure from each cluster was then selected based on a priority score defined as 1/*Resolution* – *R_free_* + *Number of protein chains*, where higher priority scores were prioritized. The final structures were visually inspected to check that they actually reflected the biological assemblies and that multiple copies of the same complex were not present in a structure (e.g. in the case of a virus capsid bound at every protein subunit by the same RNA hairpin). Final PDB IDs are listed in Supporting Information.

### Development of the Rosetta low-resolution RNP score function

Distances between RNA and protein atoms for each of the structures in the non-redundant set were calculated in Rosetta. Distributions were analyzed (below) to inform the final pairwise score terms. The RNA-protein score function is a linear combination of the previously described RNA score terms and the five RNP score terms described here. Weights for the RNP score terms were adjusted so that the magnitudes of the final score values were similar. The final score function is available within Rosetta as “rna_lores_with_rnp_aug.wts”.

### Scoring RNA-protein interactions in the plane of the base (rnp_base_pair)

A coordinate system was set up on each base as described previously (Das and Baker, 2007). Distributions of each of the protein sidechain centroids around each of the four RNA bases were analyzed for 0 < |z| < 3 Å, 0 < |x| < 10 Å, and 0 < ly| < 10 Å, and binned into 2 Å by 2 Å boxes in the x-y plane. The resulting two-dimensional histograms were smoothed with a 1.2 Å Gaussian filter (using the gaussian_filter function in python (scipy)) and normalized by the total counts of interactions across all protein sidechains and RNA bases in a given bin. rnp_base_pair scores were calculated as the negative logarithm of these frequencies. The scores were renormalized by subtracting the maximum score value across all bins, so that the maximum value of this score was equal to zero.

### Scoring RNA-protein stacking interactions (rnp_stack)

Distributions of protein sidechain centroids around each of the four RNA bases were analyzed as described for rnp_base_pair above, but above and below the plane of the base, for 3.0 < |z| < 6.5 Å. The distributions showed clear stacking interactions between each of the four RNA bases and tryptophan, tyrosine, phenylalanine, valine, and leucine. The rnp_stack score term rewards these interactions by applying a bonus of −1.0 when any of these five protein sidechain centroids falls within 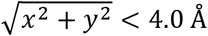 and 3.0 < |z| < 6.5 Å of an RNA base.

### Scoring general RNA-protein interactions (rnp_pair_dist)

Distances between RNA base centroids and P, C5’, C1’, C3’ atoms and protein sidechain centroids and N, C, CA, O, C atoms were binned from 0 to 20 Å in 2 Å intervals. rnp_pair_dist scores were calculated as described previously for protein-protein docking (Gray et al., 2003) (S_pair_), but because the potential here is a function of distance rather than binary interaction, the potential was normalized for interaction volume. This normalization was performed for each distance range by dividing by the total number of interactions between any protein atom and any RNA atom within that distance. Scores were adjusted so that the maximum score value was equal to zero, and to reduce noise, resulting values between 0 and −1.0 were set to 0.

### Scoring interactions with the RNA backbone (rnp_aa_to_rna_backbone)

Distances between RNA phosphate atoms and protein sidechain centroids were binned for distances between 3 and 10 Å in 1 Å intervals. Counts in each bin were normalized by the volume of the bin and by the counts of any protein sidechain in a given distance bin. The rnp_aa_to_rna_backbone scores were calculated as the negative logarithm of these frequencies. Scores were normalized to zero at 10 Å.

### Scoring steric clashes (rnp_vdw)

Interactions between each of nine RNA atoms (P, C5’, C1’, C3’, N1, and for adenosine N6, N7, N3, O2’; for cytidine N4, C6, O2, C2’; for guanosine O6, N7, N2, O2’; for uridine O4, C6, O2, C2’) and protein sidechain centroids, C, CA, O, N, and CB atoms are penalized when they come within the minimum distance observed in the non-redundant set of crystal structures. Like for the RNA and protein steric penalties (Das and Baker, 2007; Simons et al., 1997), the rnp_vdw score is calculated as 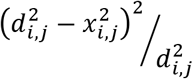 where x_i,j_ is the distance between the RNA atom i and protein atom j, and d_i,j_ is the minimum observed distance.

### Simultaneous folding and docking in Rosetta

To create *RNP-denovo*, the FARNA method was updated to include protein binding. Specifically, rigid-body docking moves are included along with the standard RNA fragment insertion moves in the Monte Carlo simulation. For runs where some RNA residues were held fixed relative to the protein (the benchmark set of ten complexes described above and telomerase described below), these rigid-body docking moves were not used, but the center of mass of the RNA could still change dramatically during modeling. Conformations are scored with the RNA-protein score function described above. This method is freely available to academic users as part of the Rosetta software package (www.rosettacommons.org). An example Rosetta command line is as follows:

~~~
rna_denovo -f fasta.txt -secstruct_file secstruct.txt -s
protein_structure.pdb -minimize_rna false -nstruct 100
~~~

where fasta.txt lists the full RNA and protein sequence, secstruct.txt contains the secondary structure in dot-bracket notation with the protein represented as dots, and protein_structure.pdb is the structure of the protein.

For each system, the RNA was built in the presence of the unbound protein structure or a homology model for telomerase and the CAPRI RNA methyltransferase. Coordinates for the fixed RNA residues for the benchmark set of ten complexes were taken from the bound complex. The orientation of the fixed RNA residues relative to the unbound protein was determined by aligning the unbound protein structure to the bound complex. For the benchmark set of ten complexes, 7000 models were generated for each system and RMSDs were calculated over the RNA heavy atoms after alignment of the protein coordinates. Further details for the three large RNA-protein systems are described below.

### Rescoring Rosetta models with docking potentials

For each of the ten systems in the benchmark set, all 7000 Rosetta *RNP-denovo* models were rescored with the DARS-RNP and 3dRPC potentials using the following commands:

For 3dRPC:

~~~
3dRPC -mode 8 -system 9 -par scoring.par
~~~

For DARS-RNP:

~~~
python ~/src/DARS-RNP_v3/DARS_potential_v3.py -f list.txt -n
~~~

### Calculating target RMSD accuracy for larger RNA-protein complexes

The size independent RMSD metric, RMSD_100_ proposed in (Carugo and Pongor, 2001) is given by:

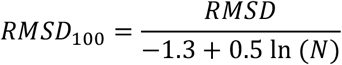

where N is the number of residues. This relationship was originally determined for proteins, but we assumed that it would also hold for RNA-protein complexes. RMSD_100_ values for the initial benchmark set and the larger RNA-protein complexes should be the same for models exhibiting the same degree of structural similarity. We then used the average number of RNA residues in the initial benchmark set (N_small_ = 21) and the larger RNA-protein complexes (N_large_ = 117), as well as the average RMSD value for the initial benchmark set (RMSD_small_ = 4.3 Å) to calculate the expected RMSD value (RMSD_large_) for the larger RNA-protein complexes:

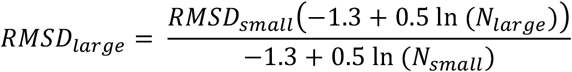

### Telomerase modeling

Models of the telomerase core RNP with an 11-nucleotide template hybrid were built without enforcing the formation of the P1 stem exactly following the procedure described previously, but using the *RNP-denovo* fold-and-dock method described here instead of the smFRET-Rosetta protocol previously described (Parks et al., 2017). As before, the template hybrid and CR4/5 RNA were kept fixed relative to the homology model of the TERT protein. Distance restraints based on FRET data were applied as described previously. As before, approximately 2500 models were built. Two additional sets of models were built following the same procedure, but without the FRET distance restraints: one with the full Rosetta RNA-protein score function and the other with only the RNA score terms and the score term penalizing RNA-protein steric clashes (rnp_vdw). RMSDs were calculated between Rosetta models and the published 7.7 Å cryoEM structure (Nguyen et al., 2018) at the ends of the pseudoknot motif over two pseudoatoms defined as the centroids of the C1’ positions of residues 96 and 118, and residues 108 and 182. RMSDs were not calculated over all atoms because the cryoEM structure was determined at 7.7 Å resolution, which is not high enough to confidently resolve positions of individual atoms. Additionally, positions of the highly conserved pseudoknot motif differ considerably between the recently determined *Tetrahymena* (Jiang et al., 2018) and human structures (Nguyen et al., 2018), and previous single-molecule FRET studies suggest conformational flexibility (Parks et al., 2017). Predicted FRET values were calculated by converting the distances between the C5 atoms in respective atom pairs to FRET with the following equation:

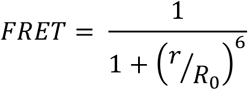

where the Förster radius, R_0_ = 56 Å and r is the distance between atoms.

### CAPRI Target 33/34 RNA methyltransferase modeling

Models of the RNA methyltransferase complex were built using either the *RNP-denovo* fold-and-dock or the prefold-then-dock method. In each case, we used the previously built protein homology model bound to the SAM cofactor (Fleishman et al., 2010) and assumed the correct RNA secondary structure. Fixed ideal A-form helices were used to model all helical elements. In both cases, constraints were applied matching those used in the original CAPRI modeling (Fleishman et al., 2010). 10000 models were built using the *RNP-denovo* fold-and-dock method. For the prefold-then-dock method, 10000 models of the RNA alone were built with FARNA. The top scoring 1000 models were clustered as described in the prefold-then-dock section above. The cluster centers of the ten most populated clusters were then docked in Rosetta to the protein homology model (using the *RNP-denovo* fold-and-dock method described here, but treating the whole RNA as a rigid body) with constraints (described above) applied. 100 models of the complex were generated for each cluster center, and the top 10 soring were taken from each to give a final pool of 100 top scoring models. RMSDs were calculated over all RNA heavy atoms after alignment over either all protein heavy atoms or all RNA heavy atoms.

### Spliceosome modeling

Models of the spliceosome core RNA bound to Prp8 were built starting from the published RNA model (Anokhina et al., 2013). This model was docked to the Prp8 crystal structure (using the same region as for the previously published model). For the *RNP-denovo* fold-and-dock runs, the position of U5 bound to the 5’ exon was allowed to move relative to the rest of the RNA model by assuming that the 3’ end of the 5’ exon is still covalently connected to the 5’ end of the intron and allowing the torsions at this connection to move freely. For the prefold-then-dock runs, the published RNA model was treated as a rigid body. The three crosslinking constraints used to build the published model of the RNA docked to Prp8 were also included here. Specifically, a score penalty was applied to conformations where none of the C1 ‘ atoms of the U5 RNA residues were within 5.0 Å of residues 1281-1414 of Prp8. An additional penalty was applied when there were no RNA branchpoint helix residues within 5 Å of Prp8 residues 1575 or 1598 (the ends of the disordered loop in Prp8). Finally, a penalty was also applied when there were no residues within the ACAGA/pre-mRNA loop within 30 Å of residue 1826 of Prp8 (the connection to the RNaseH-like domain, which was not included in the models, and is meant here to represent its approximate location). All of these restraints were implemented as ambiguous flat harmonic restraints in Rosetta with standard deviations of 1 Å. Because these restraints were so stringent, they were considered “satisfied” in the final models when the total restraint score was less than 60.0 Rosetta energy units and as an additional check, U5 residue 39 was within 60 Å of the center of mass of Prp8 residues 1281-1323+1326-1413, the fourth intron residue in the branchpoint helix was within 25 Å of the center of mass of Prp8 residues 1575 and 1598, and intron residue U+4 was within 60 Å of Prp8 residue 1826. Approximately 1500 models were built. RMSDs were calculated over phosphorus atoms in the last (3’) residue of the 5’ exon, the first residue (5’) of the intron, and the residue right after (3’ of) the branchpoint A with respect to the corresponding residues from the published cryoEM structure of the yeast C complex (Wan et al., 2016) after alignment over Prp8.

